# Diet composition, body condition and sexual size dimorphism of the common African toad *(Amietophrynus regularis)* in urban and agricultural landscape

**DOI:** 10.1101/2021.01.25.428067

**Authors:** Benjamin Yeboah Ofori, John Bosu Mensah, Roger Sigismund Anderson, Daniel Korley Attuquayefio

## Abstract

Land use and land cover change (LULCC) are major drivers of global biodiversity loss. The conversion of natural habitats into human-modified landscapes poses novel and multifaceted environmental stressors to organisms, influencing their ecology, physiology, life history and fitness. Although the effects of LULCC have been studied extensively at the community level, there is scant information about its effect on population and individual characteristics. We assessed the diet composition, body condition, and sexual size dimorphism of the common African toad *(Amietophrynus regularis)* in urban and agricultural landscape. Diet composition was evaluated using gut content analysis, while body condition was measured using residual mass index. Overall, 935 prey items comprising six classes, at least 18 orders and 31 families were obtained from toads. This broad dietary niche suggested that *Amietophrynus regularis* is a generalist predator. The family Formicidae was the most consumed prey item, with a frequency of occurrence above 80% at both sites. We found no sex- or habitat-biased dietary partitioning in the toads. A statistically significant positive correlation existed between snout-vent-length (SVL) and diversity of prey items (Pearson’s correlation r = 0.999, p ≤ 0.0001) for toads from farmland, which also had better body conditions. The toads showed female-biased sexual size dimorphism, but males had longer tibio-fibula, radio-ulna, foot, and distal fore limbs. This study is probably the first to assess the diet composition, body condition and sexual size dimorphism of *Amietophrynus regularis* simultaneously. The ecological, evolutionary and conservation implications of our findings are discussed.

## Introduction

Agricultural intensification and urbanization are occurring at an alarming rate globally, leading to land cover changes and simplification of natural environments (Mullu, 2016; Carraso *et al*., 2017). Land use and land cover changes are recognized as major drivers of global biodiversity loss (Concepcion *et al*., 2015; Lindenmayer, 2019; Liu *et al*., 2018). The conversion of forests and other natural vegetation into human settlements and agricultural landscapes poses multifaceted challenges and novel environmental stressors that threaten the survival of local species (Wilson *et al*., 2016; Rakotoniana *et al*., 2016; Salomao *et al*., 2018). Organisms inhabiting small habitat fragments may experience high levels of environmental stressors, such as climate change, increased competition, predation, parasitism, and decreased food availability (Mikolas, 2016). This may cause poor body condition, reduced fecundity, genetic diversity, reproductive success and fitness of organisms (Janin *et al*., 2011), which ultimately reduce the diversity and abundance of fauna and flora species and cause alterations in community composition and biotic interactions (Newbold *et al*., 2015; Segan *et al*., 2016; Fletcher, 2018; Liu *et al*., 2018; Jiménez-Peñuela *et al*., 2019).

The characterization of the ecological niche of a given species is based on its distinct life history, particularly its use of food resources as one of the most studied by ecologists (Moser et al., 2017). Food is a fundamental resource that influences organisms in a variety of ways (Marshal et al., 2009). The pursuit and consumption of food affects the behavior, ecology, and life history of organisms (Lee-Thorp and Sponheimer, 2006). The ability of an individual to safely harvest and process sufficient quality food to fulfill its requirements for growth, maintenance, and reproduction is a key determinant of its fitness (Marshal et al., 2009). Knowledge of the diet of an organism and its changes in space and time is therefore essential for understanding its ecological role and population fluctuations (Ifarraguerri *et al*., 2017).

The stress imposed by LULCC on organisms may affect their nutritional status, energetic state and physiology, ultimately causing declines reproductive rates and populations (Mikolas, 2016). Body condition has been used extensively as a measure of environmental stress imposed on organisms, such as intense competition, predation and parasitism (Green, 2001; Resano-Mayor *et al*., 2016; Ronto and Rakotondravony, 2019). It has also been used as a measure of habitat quality, foraging success, reproductive investment, mate choice, fighting ability and disease susceptibility (Jarvis, 2015; Falk *et al*., 2017; Bancila *et al*., 2019). Body condition reflects the physiological state, energetic status, health and fitness of organisms (Cox and Calbeek, 2015; Resona-Mayor *et al*., 2016; Warner *et al*., 2017). Individuals with better body condition are expected to possess good health and greater energetic status, and consequently better chances of survival than those with poorer body condition (MacCracken and Stebbings, 2012; Labocha *et al*., 2014).

Sexual dimorphism refers to the existence of intraspecific differences in morphology, coloration, ornaments and body size between males and females (Kupfer, 2009; Altunisik, 2017; Mori *et al.*, 2017). It is widespread among animals (Seglie *et al*., 2010; Di Cerbo and Biancardi, 2012; Ivanovic and Kalevic, 2012; Labus *et al*., 2013), varying widely even within closely-related groups (Radojicic *et al*., 2002; Paoletti *et al*., 2009; De Lisle and Rowe, 2015). Sexual dimorphism arises from different selective pressures, such as natural selection, sexual selection and fecundity selection (Vargas-Salinas, 2006; Kupfer, 2009; Liao *et al*., 2012; Martin *et al*., 2012; Colleoni *et al,* 2014; Altunisik, 2017). Natural selection theory posits sexual dimorphism as due to differential interactions of each sex with the environment (Lovich and Gibbons, 1992). Natural selection acts on traits that favor larger males or females depending on the ecological context (Colleoni *et al*., 2014).

According to sexual selection theory, sexual dimorphism arises from competition for mates and is attributed to sexual differences in reproductive roles (Seglie *et al*., 2010). Traits such as bright coloration, hypertrophied morphological features or large body size that enhance access of one sex to the opposite sex, are favoured (Lovich and Gibbons, 1992). Larger males are favoured, and gain an advantage in intrasexual competition with other males (Martin *et al*., 2012; Zhang and Lu, 2013; Colleoni *et al*., 2014; Magalhaes *et al*., 2016; Altunisik, 2017). Sexual selection may also lead to dietary differences between sexes (Magalhaes *et al*, 2016). Unlike sexual selection, fecundity selection favors larger females, possibly promoting larger clutch sizes and enhanced reproductive capacity (Lowe and Hero, 2012; Martin et al., 2012; Zhang and Lu, 2013; Colleoni *et al*., 2014). For instance, females are known to be larger than males in 90% of anurans and 61% of caudates (Marzona *et al*., 2004; Seglie *et al*., 2010; Zhang and Lu, 2013; Altunisik, 2017).

Amphibians are among the most species-rich groups of terrestrial vertebrates, with hundreds of new species still being discovered annually (Musah *et al*., 2019). They are important components of ecosystem, influencing food webs and energy flow by their feeding behaviour (Le *et al*., 2018) and by serving as prey for larger vertebrates (Park *et al*., 2018). Despite their ecological importance, diversity and high vulnerability, Afro-tropical amphibians have received relatively less attention in the ecology, evolution and conservation literature. Most of the studies on amphibians are conducted in the Neotropics and concentrate on their distribution and speciation. There is scant information on amphibian population dynamics and the influence of land cover/land use changes on their ecology and life-history traits.

Most amphibians are generalist and opportunistic invertebrate feeders (Balint *et al*., 2008; Crnobrnja-Isailovic *et al*., 2012; Kittel and Sole, 2015; Do Couto *et al*., 2018; Santana *et al*., 2019). However, some species are specialist predators (Menin *et al*., 2015; Do Couto *et al*., 2018). The feeding habit and diet of amphibians may be influenced by seasonal availability and abundance of food (Crnobrnja-Isailovic *et al*., 2012), presence of competitors, and predation risk (Lopez *et al*., 2009; Kittel and Sole, 2015; Santana *et al*., 2019). Therefore, the ability of an individual amphibian to safely harvest and process sufficient quality food may vary with environmental differences. This variation may be reflected in the diet composition and body condition of individuals. Also, individuals living in environments with high competition and risk of predation may develop adaptive traits that may differentiate them from individuals in less stressful environments.

The common African toad *(Amietophrynus regularis)* is common in Sub-Saharan Africa (Vancocelos *et al*., 2010; Iyaji *et al*., 2015). However, little is known about its ecology and life history as well as its trophic niche and how this is influenced by LULCC. Currently, there is lack of knowledge on the extent to which urban and agricultural landscapes influence their diet composition and body condition, as well as their adaptations for fitness enhancement under these stressful environments. Such information is of ecological and evolutionary significance and also important for predicting the impact of environmental changes on individuals and populations and for formulating effective conservation strategies.

The present study therefore assessed the diet composition, body condition and sexual dimorphism of the common African toad from agricultural and urban landscapes in Accra, Ghana. We hypothesized that agricultural and urban landscapes may have different effects on the diet composition, body condition and prominence of sexual size dimorphism of the common African toad. We expected toads from farmland to have more diverse diet and better body condition than those from the urbanized areas. Like most anurans, we expected females to have bigger body size than males. We also expected males from the urban area to have prominent traits that enhance sexual selection because of intense competition for mates.

## Materials and Methods

### Study Area

This study was conducted on Legon campus (05°39’03”N, 00°11’13”W) of the University of Ghana. The campus has a total area of 1,300 hectares (13 km^2^) and is located about 13 km north-east of Accra, the capital city of Ghana (Gbogbo *et al*., 2017). The climate of the area is characterized by a pronounced gradient of mean annual rainfall ranging from 733 mm to 1,118 mm distributed over a major (May-July) and minor (September-October) rainy season and a daily mean temperature of about 30°C (Holbech *et al*., 2015; Gbogbo *et al*., 2017). The vegetation is generally coastal grassland, thicket and dry forest (Garshong *et al*., 2013), with most of the natural vegetation converted into developed areas (buildings, markets, roads and other man-made infrastructure) and farmland. The only remnant of the original vegetation is within the Legon Botanical Garden which is located north of the campus and covers an area of about 2.0 km^2^. The developed (urban) area, which covers about 7.3 km^2^, includes areas around students’ halls, hostels, chalets, staff bungalows, academic facilities, central administration blocks, library block, bookshops, banking square, restaurants and canteens. The farmland is located adjacent to the botanical garden and covers an area of about 1 km^2^. Crops grown in the farm include mango, cashew, maize and vegetables.

### Study Species

The common African toad, *Amietophrynus regularis [Bufo regularis]* (Amphibia: Bufonidae) (Iyaji *et al*., 2015), is a large, strong toad with warty skin and dark olive-brown dorsal surface. The underside of both sexes is white to beige, with males having a black sub-gular or throat pigmentation (Deef, 2019). The common African toad also known as the square-marked toad, is listed as ‘Least Concern’ by the IUCN in view of its wide distribution in a broad range of habitats and presumed large population size (IUCN, 2016). This toad occurs in both moist and dry savanna, forest margins, montane grassland, and agricultural and other habitats associated with rivers (Deef, 2019). It is widely distributed in Sub-Saharan Africa, with its range extending from oases in Algeria and Libya to northern Nilotic Egypt (Ibrahim, 2008; Saber *et al*., 2016). The African common toad feeds on a wide variety of vertebrates and invertebrates, particularly insects (Ibrahim, 2008), serving as insect biocontrol in most agricultural fields (Saber *et al*., 2016). It is a good source of protein in many parts of sub-Saharan Africa, notably Nigeria and Burkina Faso (Akinsanya *et al*., 2020).

### Data Collection

Using visual encounter surveys, samples were collected from farmland and developed areas on the University of Ghana, Legon campus. Prior to sampling, the study area was surveyed and breeding sites determined using points of male calls. Samples were collected from July to September between the hours of 22:00GMT and 08:30GMT. During the day, toads were searched for under leaf litter, fallen logs, rocks, and other microhabitats where toads are known to seek refuge. During chorusing nights, calling males or pairs in amplexus (Plate 1) were captured by hand following Quiroga *et al*., (2015) and Oropeza-Sanchez *et al.* (2018). The sexes of individuals were determined using the presence or absence of sub-gular pigmentation (Plate 2) following Vera-Candioti *et al.* (2019).

**Plate 1:**
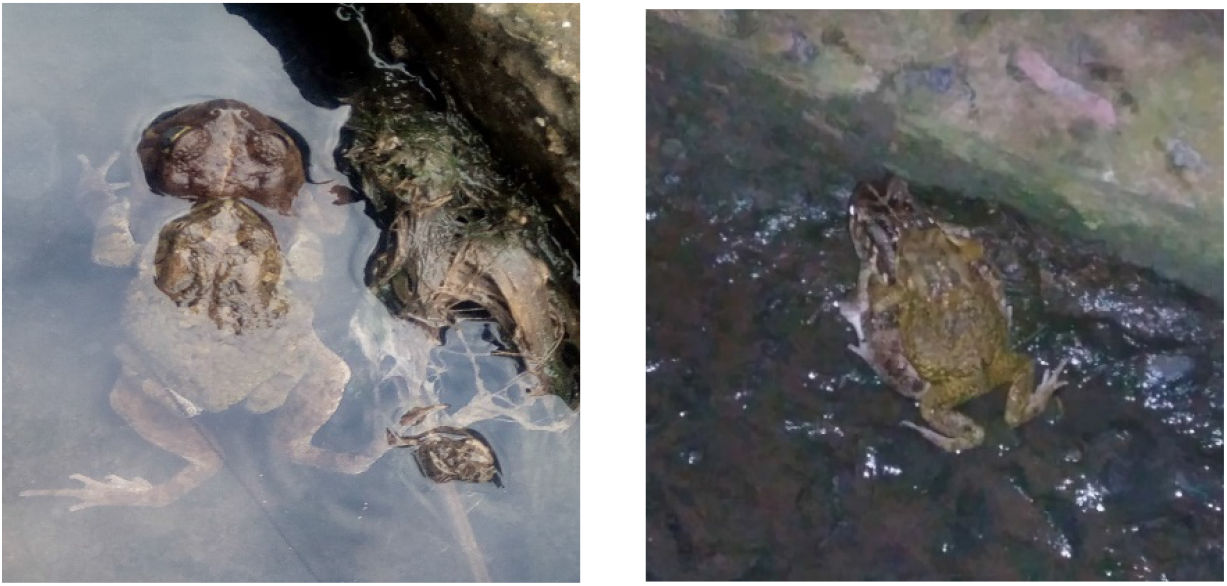
Toad pairs in amplexus encountered at night in the Developed Area

**Plate 2:**
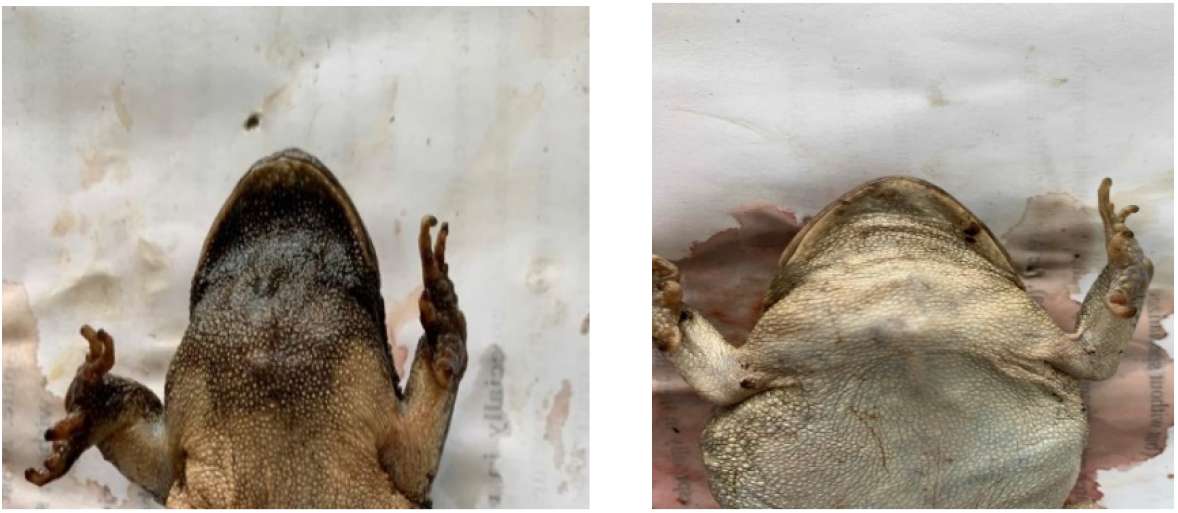
Differentiation of males from females through sub-gular pigmentation, which is darker in males.

Morphometric parameters including snout vent length (SVL), weight (W), mouth width (MW), head width (HW), head height (HH), head length (HL), foot length (FT), radio-ulna length (RUL), humerus length (HuL), femur length (FL) and tibia-fibula length (TbFL) were measured for each captured toad (Table 1). The diet composition of toads was assessed using gut content analysis. The toads were dissected and stomach contents were emptied into a petri dish containing 70% ethanol. Identification of stomach content was done to the highest taxonomic level possible (order or family) using a dissecting microscope.

**Table 1:** Morphometric measures and their respective explanations.

| **Measured Parameter** | **Abbreviation** | **Explanation** |
| --- | --- | --- |
| Snout Vent Length | SVL | Distance from the tip of the snout to the extreme posterior point of the cloaca. |
| Head Length | HL | Distance between the tip of the snout and the posterior side of the head. |
| Head Width | HW | Broadest distance between the external sides of the head. |
| Head Height | HH | Distance from the highest portion of the head to the bottom of the lower jaw. |
| Mouth Width | MW | Distance between both points of the mouth. |
| Humerus Length | HL | Distance from the posterior margin of the fore limb (axilla) to the elbow. |
| Radio Ulna Length | RuL | Distance from the elbow to the wrist of the forelimb. |
| Distal Fore Limb | DFL | Distance from the wrist to the tip of the longest digit of the forelimb. |
| Femur Length | FL | Linear distance from the ischium to the knee. |
| Tibio Fibula Length | TbBL | Linear distance from the knee to the tarsal articulation. |
| Distal Hind Limb | DHL | Linear distance from the knee to the tip of the longest digit of the hind limb. |
| Hind Foot | FT | Linear distance from the heel to the tip of the longest digit of the hind limb. |
| Weight | WT | Mass of toad. |

### Data Analysis

We calculated the Numerical Percentage (N%) as (N*i* x 100)/N*t*, where Ni is the number of prey category *i,* and N*t* is the total number of all prey categories. The Frequency of Occurrence (F%) was estimated as the number of toad stomachs in which category *i* prey were found. Diversity of Diet Composition was estimated using the Shannon-Weiner index *H’* as -∑ [(*pi*) ×ln (*pi*)], where *‘pi* represents the proportion of total abundance represented by the *i*^th^ species. The Sorenson’s qualitative index and Bray-Curtis quantitative index were used to assess the similarity of diet composition between sexes and sites. The Pearson’s correlation coefficient (*r*) was used to assess the correlation between SVL and diversity and abundance of prey categories consumed by the toads. The residual mass index was calculated using log_10_ transformed data to insure the linearity of the relationship between weight (W) and snout-vent length (SVL). The theoretical body weight of each frog was obtained by introducing the SVL into the equation of the regression line. This was then subtracted from the measured body weights to obtain the residual value (Bacila *et al*., 2010; Vera-Candioti *et al*., 2019). Individuals with positive residual indices were considered to be in good condition, while those with negative residual indices were said to be in poor condition. The body condition scores were compared between sites and sexes, and the degree of significance evaluated using the t-test. The t-test was also used to assess the significance of the differences between morphometric parameters of males and females from farmland and developed areas. A trait was said to be female-biased if the male to female ratio was less than 1, and male-biased if the ratio is greater than 1. No sexual size dimorphism existed where the male to female ratio equalled 1. All analyses were in Excel and SPPSS version 22 and test of significance was at alpha of 0.05.

## Results

### Diet Composition

A total of 44 (21 males and 23 females) and 48 (27 males and 21 females) individual toads were captured from the Farmland and Developed Area, respectively. Thirty-six percent of the Farmland individuals had empty stomachs, while 26.2% of the Developed Area individuals had empty stomachs. The compositions of diet consumed by toads were highly similar (Sorenson’s index = 83.9%; Bray-Curtis index = 91.6%), but the diversity was higher in the Farmland (Shannon-Weiner index *H’* = 1.885) than Developed Area *(H’* = 1.081). A total of 428 prey items, consisting of six invertebrate classes (Insecta, Arachnida, Chilopoda, Diplopoda, Gastropoda, and Polychaeta) belonging to at least 16 orders and 31 families were counted from the Farmland. Majority (88.8%) of these were arthropods of which 82.5% were insects. The most abundant prey items were from the order hymenoptera (39.5%), Coleoptera (15.7%), and Diptera (12.9%). Formicidae and Stratiomyidae were the most abundant insect families, with 38.6% and 11.4% of prey items, respectively. Of the non-insect orders, Polychaetes were the most frequent prey category, occurring in 50% of individuals with gut content, followed by arachnids (25%) and Chilopods (21.4%) (Table 2). For the class Insecta, the most frequent prey category was the Formicidae (ants), followed by Carabidae (39.3%) and Gryllidae (35.7%) (Table 2).

**Table 2:**
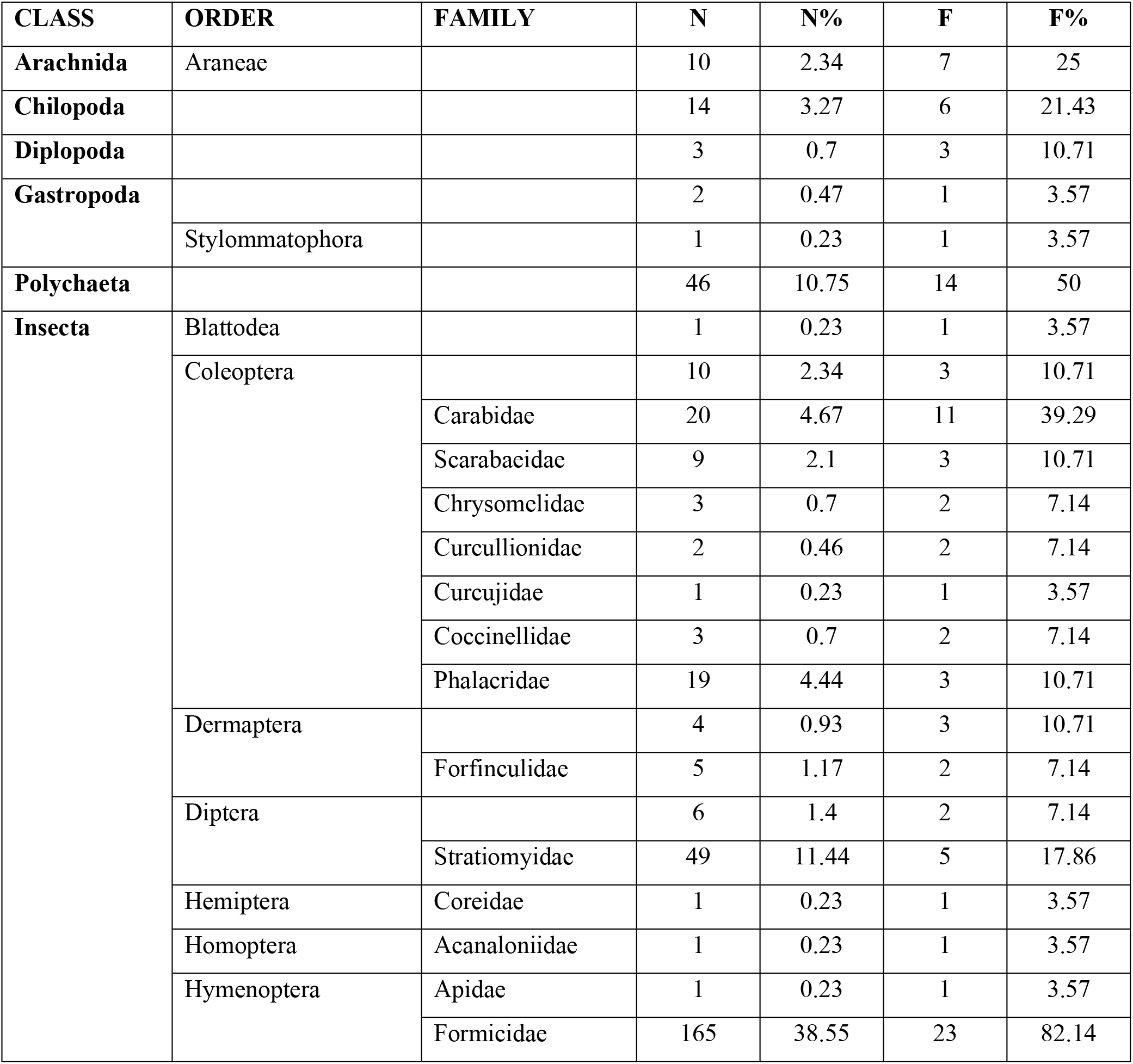
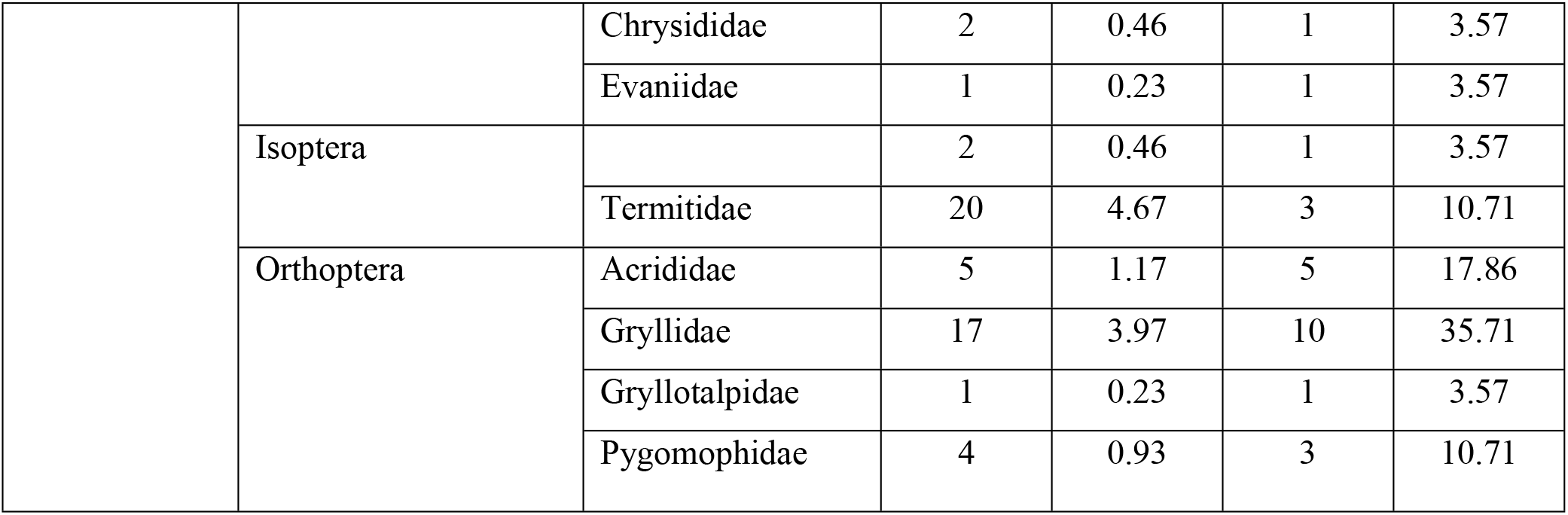
Taxonomic composition of prey items (N = 428) found in stomachs of *Amietophrynus regularis* from the University Farm. (**N =** Number of items, **N% =** Numerical percentage, **F** = frequency of occurrence and **F%** = percentage frequency of occurrence).

Females consumed a higher diversity *(H’* = 1.905) of prey items than males *(H’* = 1.627). At least, 15 and nine orders of prey items were consumed by females and males respectively from the Farmland, with all the orders of prey items found in males also being present in females (Fig. 2). The similarity (Sorenson’s index) of prey items in terms of order was therefore very high (75%). A very strong statistically significant positive correlation (Pearson’s correlation, *r* ≥ 0.9996, p < 0.00001) existed between SVL and diversity of prey items consumed for both males and females (Fig. 4). There was also a weak positive correlation (Pearson’s correlation *r* ≤ 0.3693, p ≥ 0.0561) between SVL and the abundance of prey items consumed (Fig. 4).

**Figure 1:**
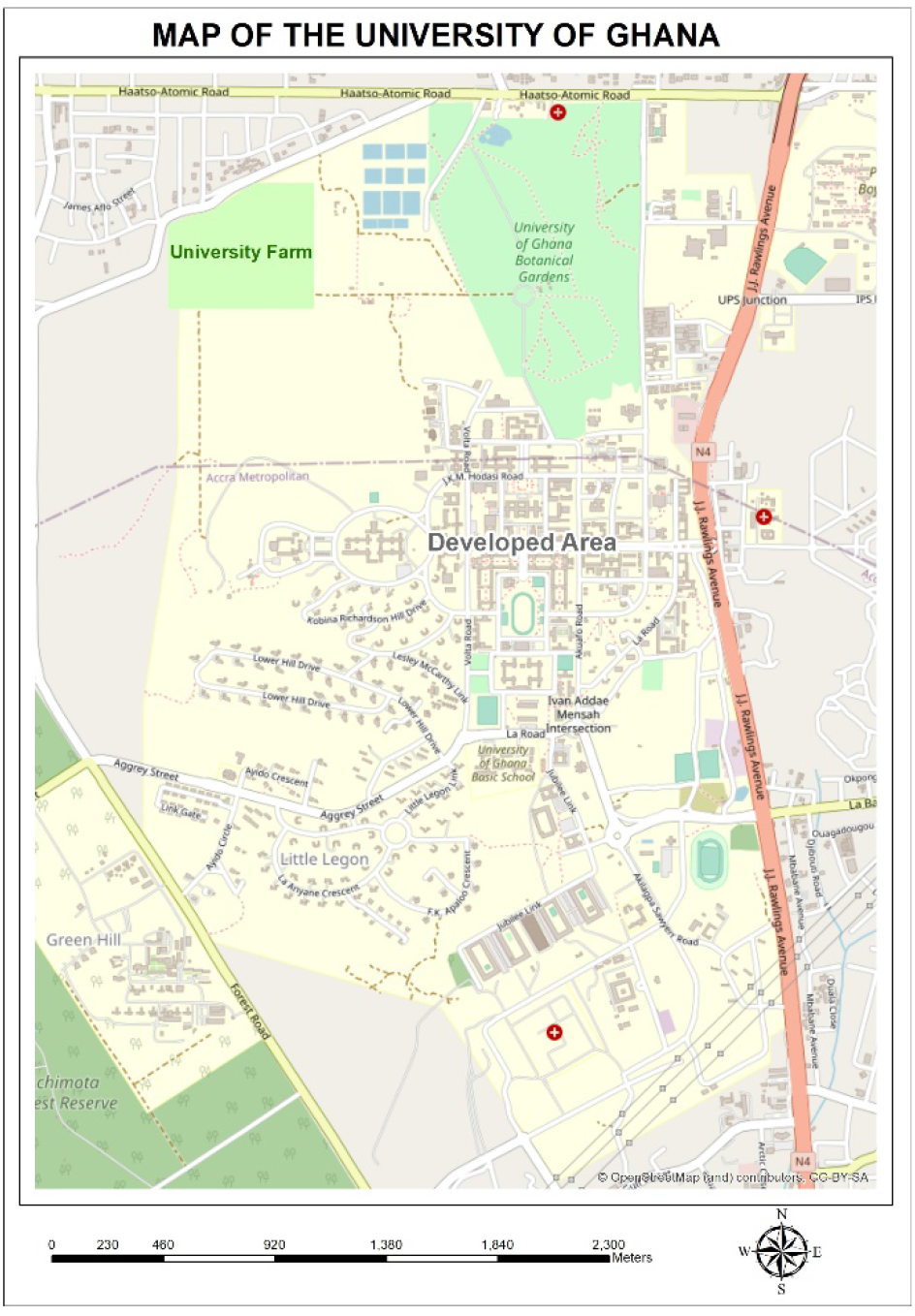
Study area showing the Developed areas and University farms

**Figure 2.**
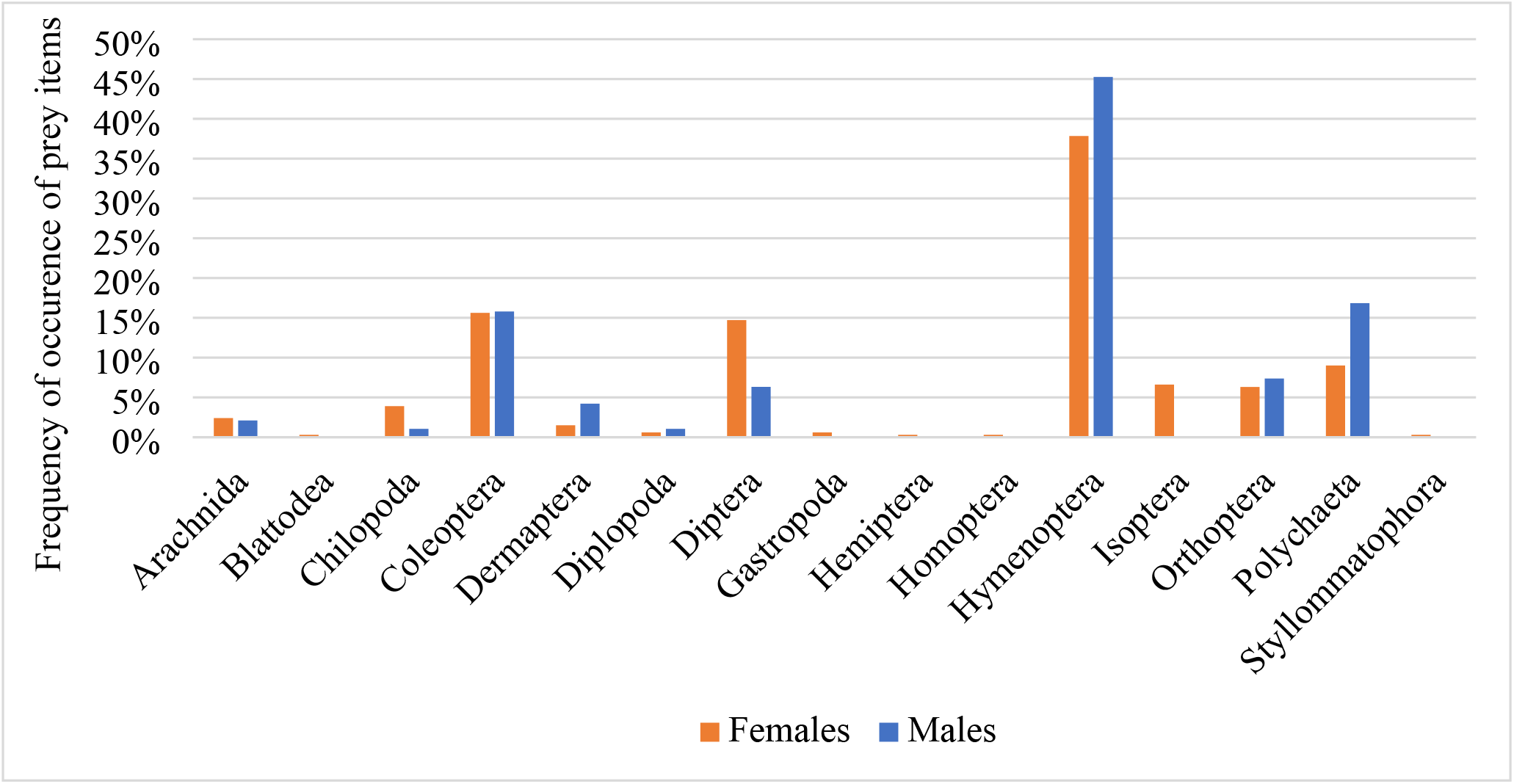
Frequency of occurrence of prey items in the gut of African common toads from the farmland.

**Figure 3.**
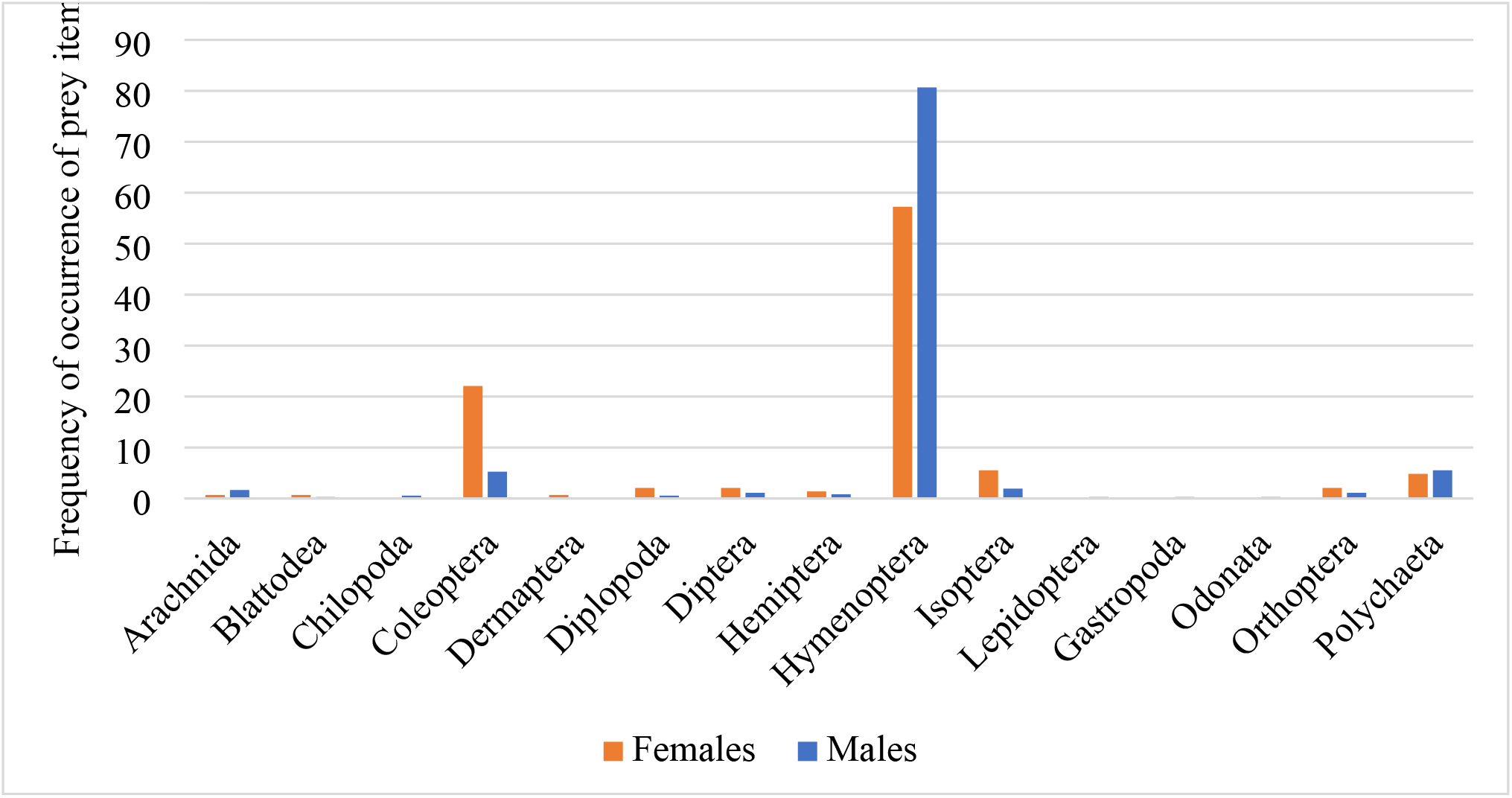
Frequency of occurrence of prey items in the gut of African common toads from the developed areas.

**Figure 4.**
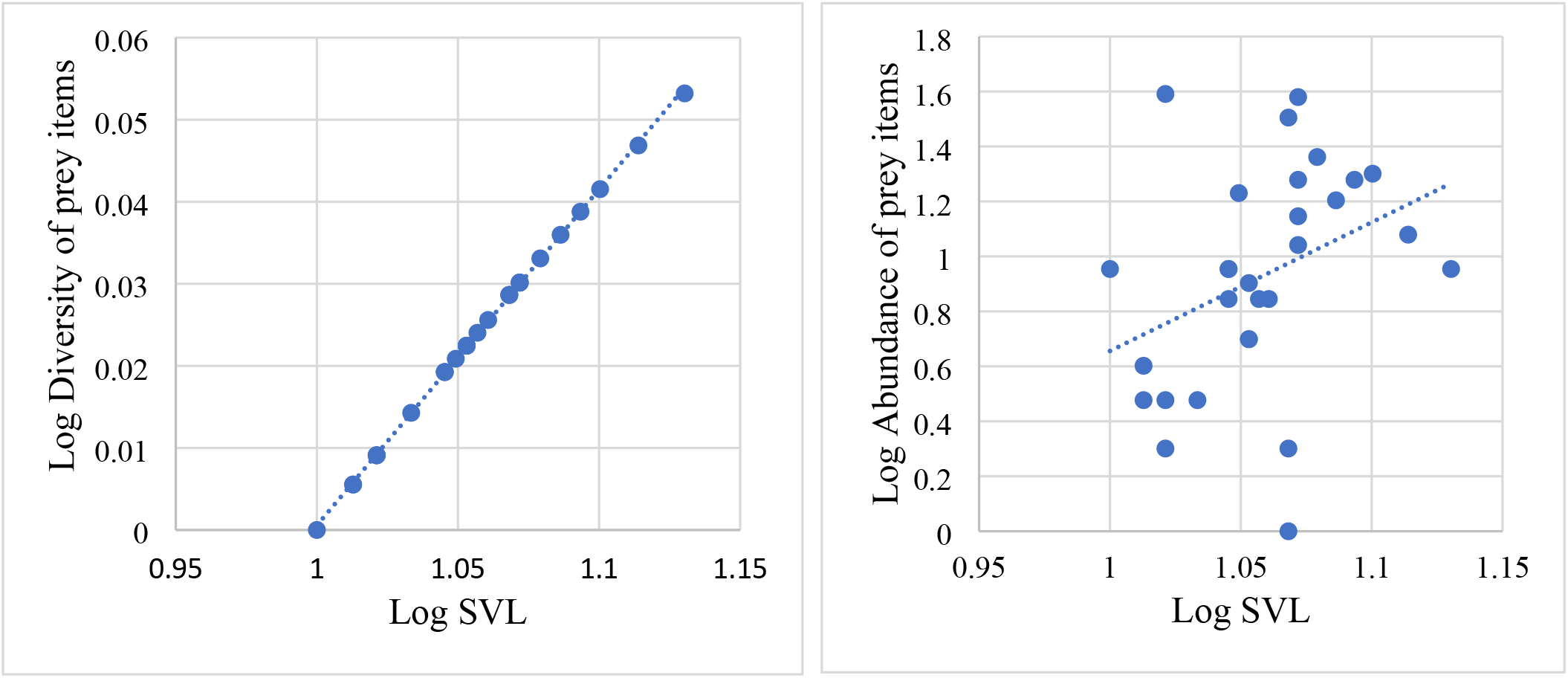
Correlation between SVL, diversity and abundance of prey items consumed by the African common toad from the farmland. (Pearson’s correlation r = 0.999, p < 0.00001; r = 0.3586, p = 0.0561).

A total of 507 prey items, comprising six invertebrate classes (Insecta, Arachnida, Chilopoda, Diplopoda, Gastropoda, and Polychaeta), from at least 15 orders and 28 families were obtained from toads in the Developed Area. The class Insecta formed the majority (91.7%) of the prey items, with the Hymenoptera (74%) and Coleoptera (10.1%) being the most abundant prey categories (Fig. 3). The Formicidae was the most abundant family (74%), and the most consumed prey category with 91.1% of individuals with gut content consuming it. Males consumed more quantities of prey items in terms of numerical proportion, but females consumed a higher diversity of prey items *(H’* = 1.364) than males *(H’ =* 0.894). There was a weak positive correlation between SVL and the diversity of prey items consumed for both males and females (Pearson correlation coefficient *r* = 0.344, p = 0.192, male; *r* = 0.1199, p = 0.536, female) (Fig. 7). There was also a weak positive correlation between SVL and the abundance of prey items consumed by females (*r* = 0.053, p = 0.846) (Fig. 8), but males showed a weak negative correlation between SVL and the abundance of prey items consumed (*r* = −0.168, p = 0.384) (Fig. 9).

**Figure 5.**
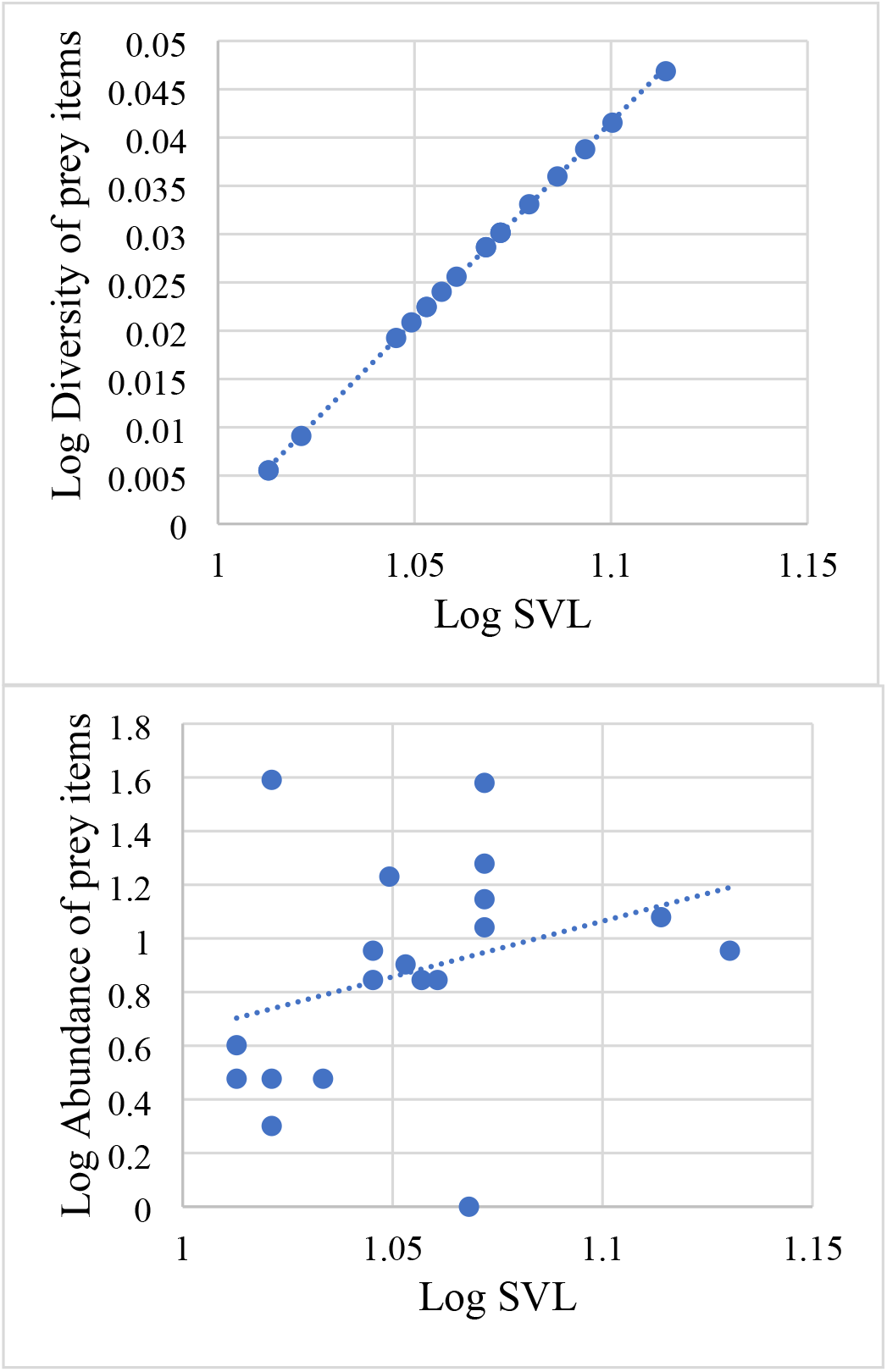
Correlation between SVL, diversity and abundance of prey items consumed by female African common toad from the farmland. (Pearson’s correlation coefficient r = 0.9999, p < 0.00001; r = 0.3693, p = 0.1197).

**Figure 6.**
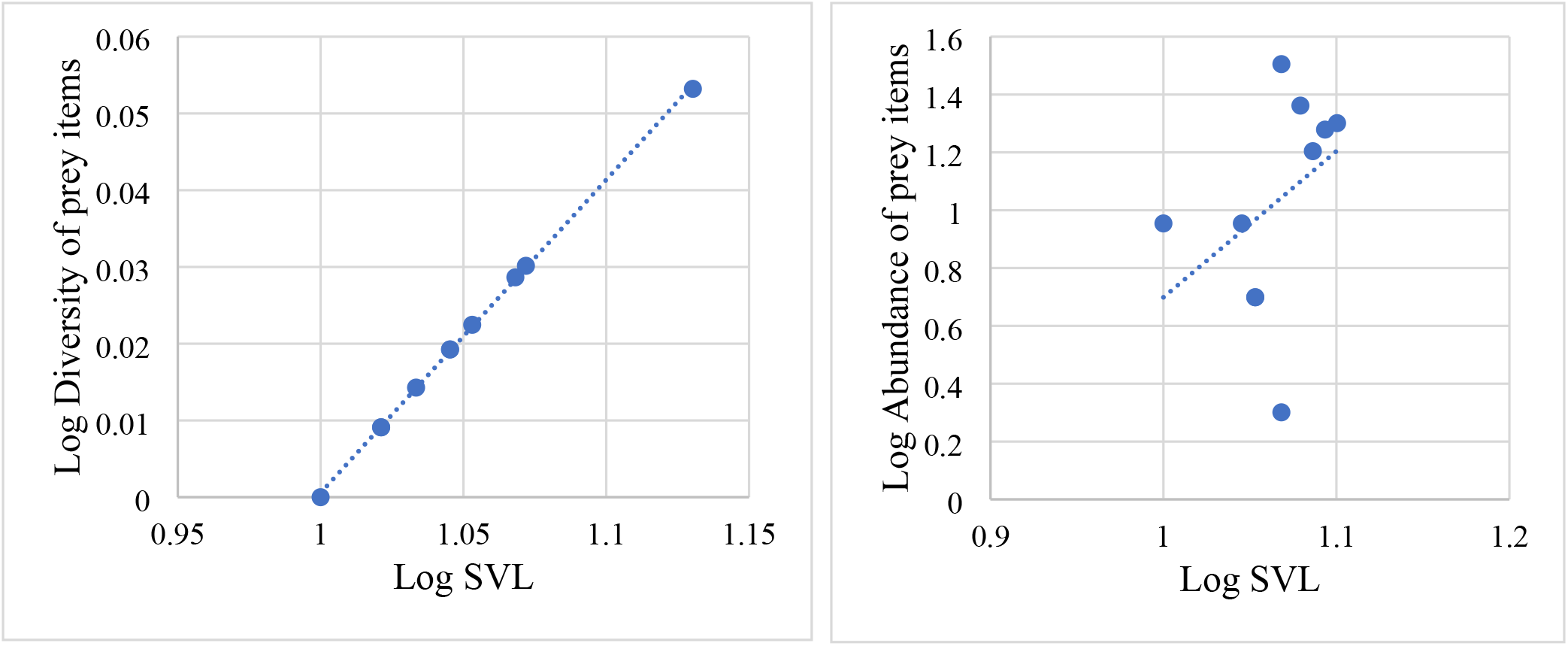
Correlation between SVL, diversity and abundance of prey items consumed by male African common toad from the farmland. (Pearson’s correlation coefficient r = 0.9998, p < 0.00001; r = 0.1952, p = 0.5889).

**Figure 7.**
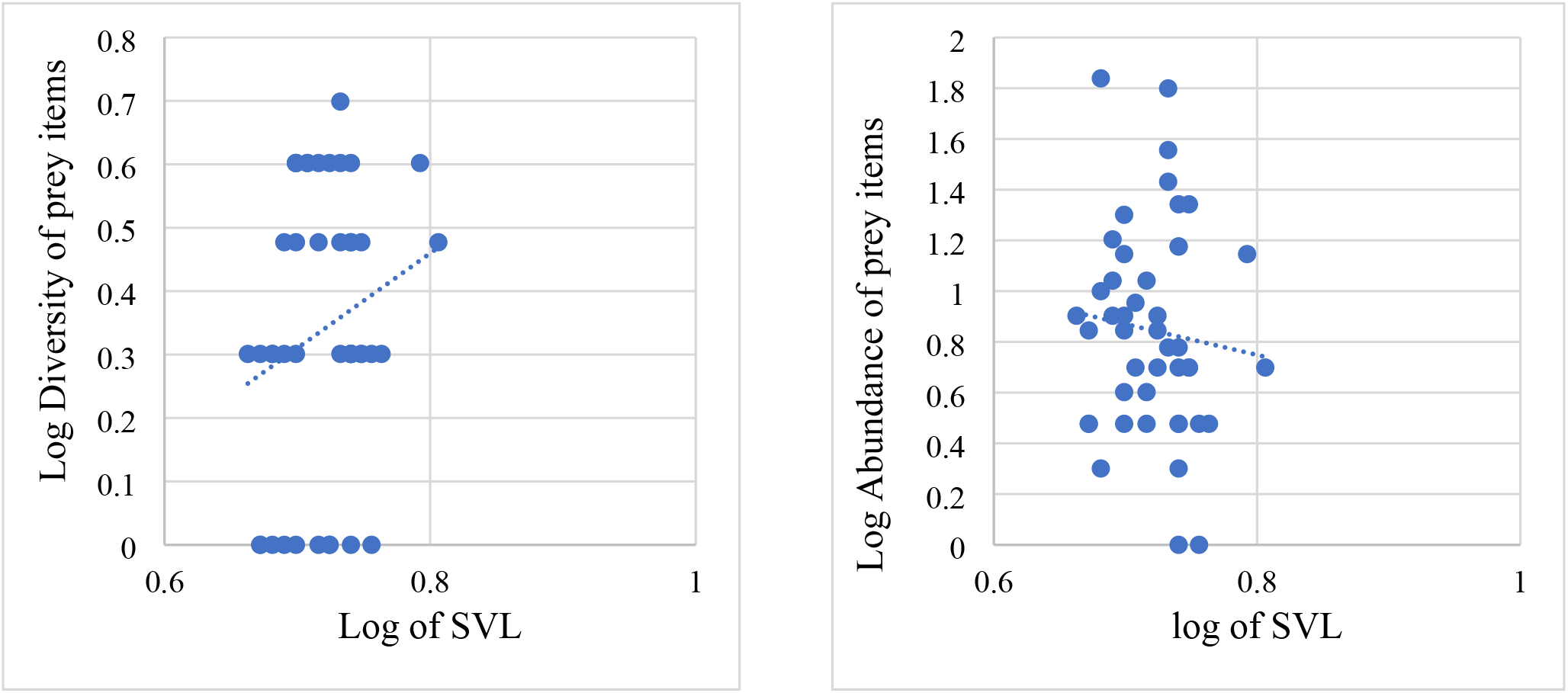
Correlation between SVL, diversity and abundance of prey items consumed by African common toads from developed areas. (Pearson’s correlation coefficient r = 0.215, p = 0.157; r = −0.094, p = 0.539)

**Figure 8.**
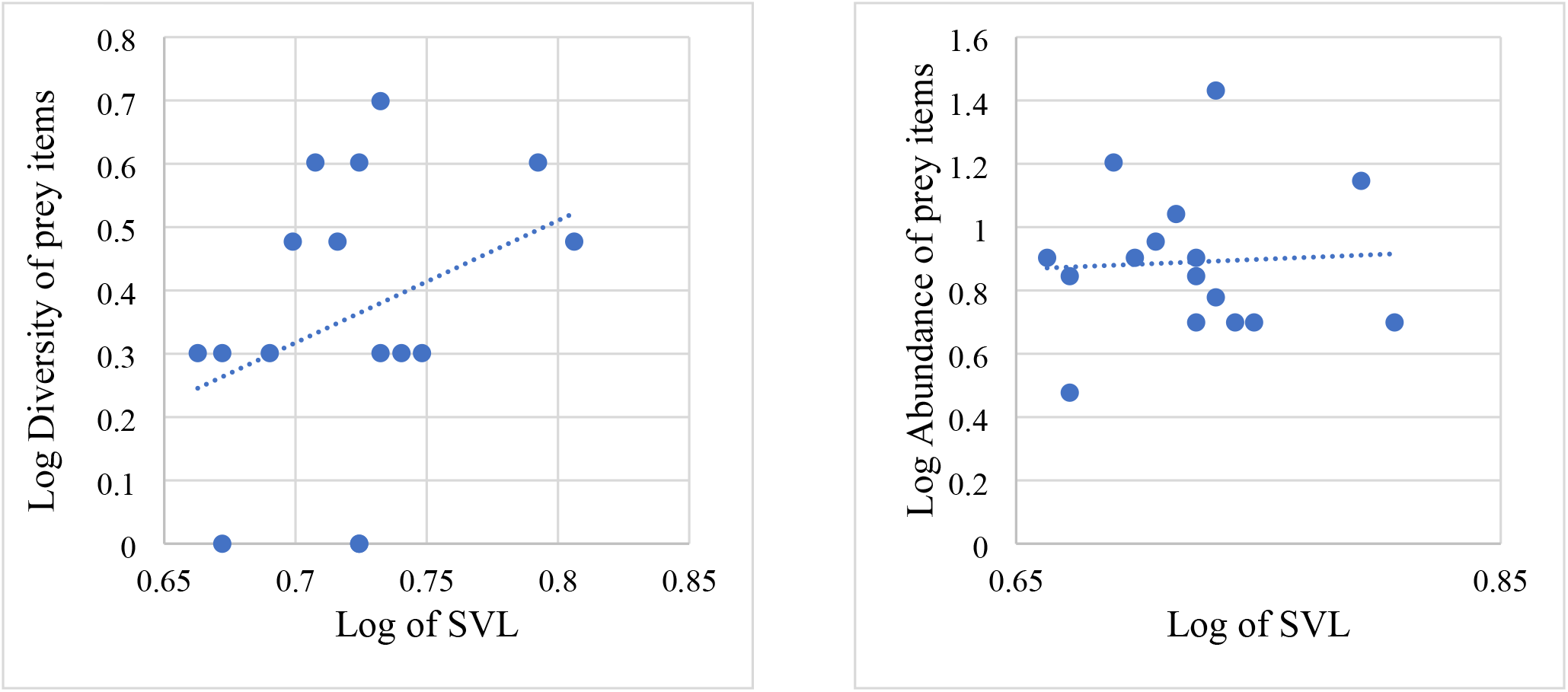
Correlation between SVL, diversity and abundance of prey items consumed by female African common toads from developed areas. (Pearson’s correlation coefficient r = 0.344, p = 0.192; r = 0.053, p = 0.846)

**Figure 9.**
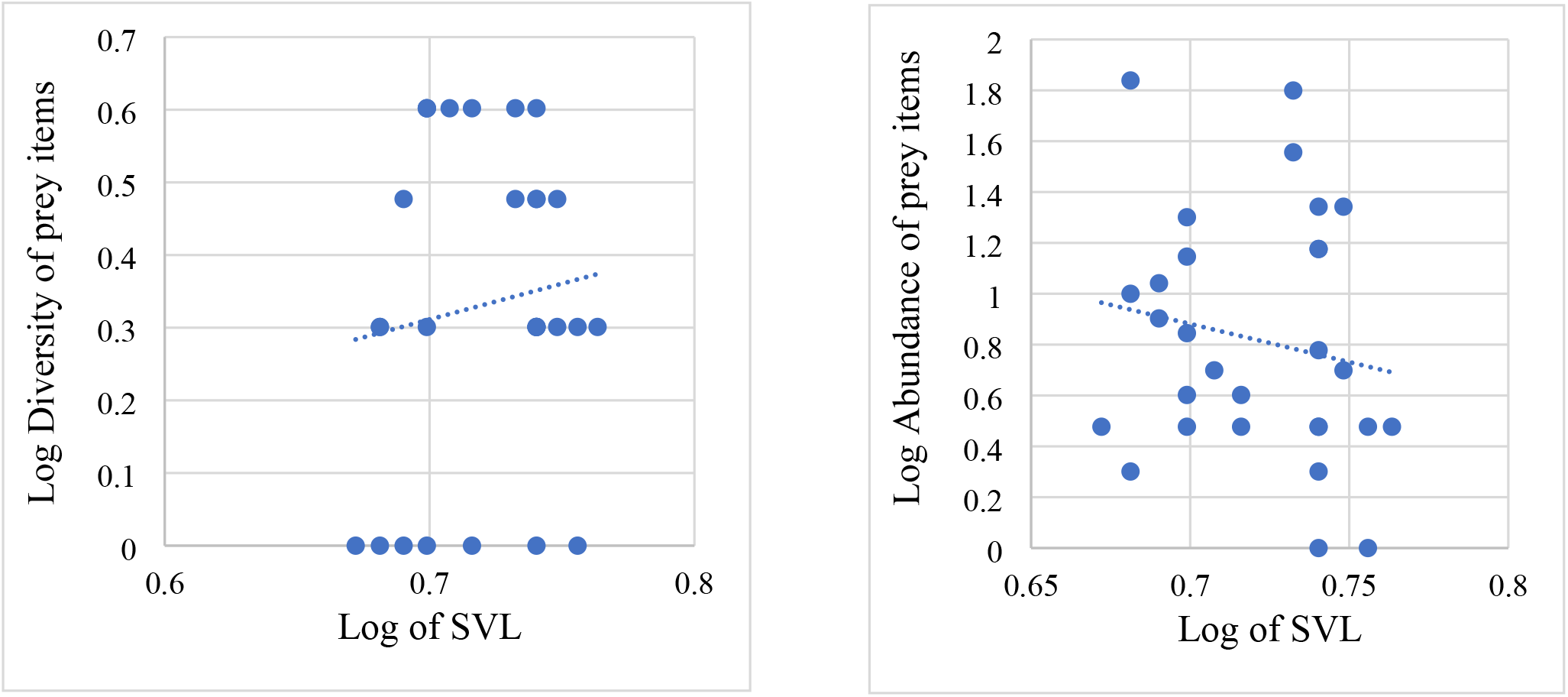
Correlation between SVL, diversity and abundance of prey items consumed by male African common toads from developed areas. (Pearson’s correlation coefficient r = 0.1199, p = 0.536; r = – 0.168, p = 0.384)

### Body Condition

Generally, toads in the Farmland and Developed Areas had poor body condition (i.e., negative residual mass index), except females from the farmland. However, both male and female individuals from the Farmland had better body condition than those from the Developed Areas. Also, male toads from the Developed Areas appeared to possess better body conditions than their female counterparts, even though the difference was not statistically significant (t-test, alpha = 0.05, p = 1).

**Figure 10.**
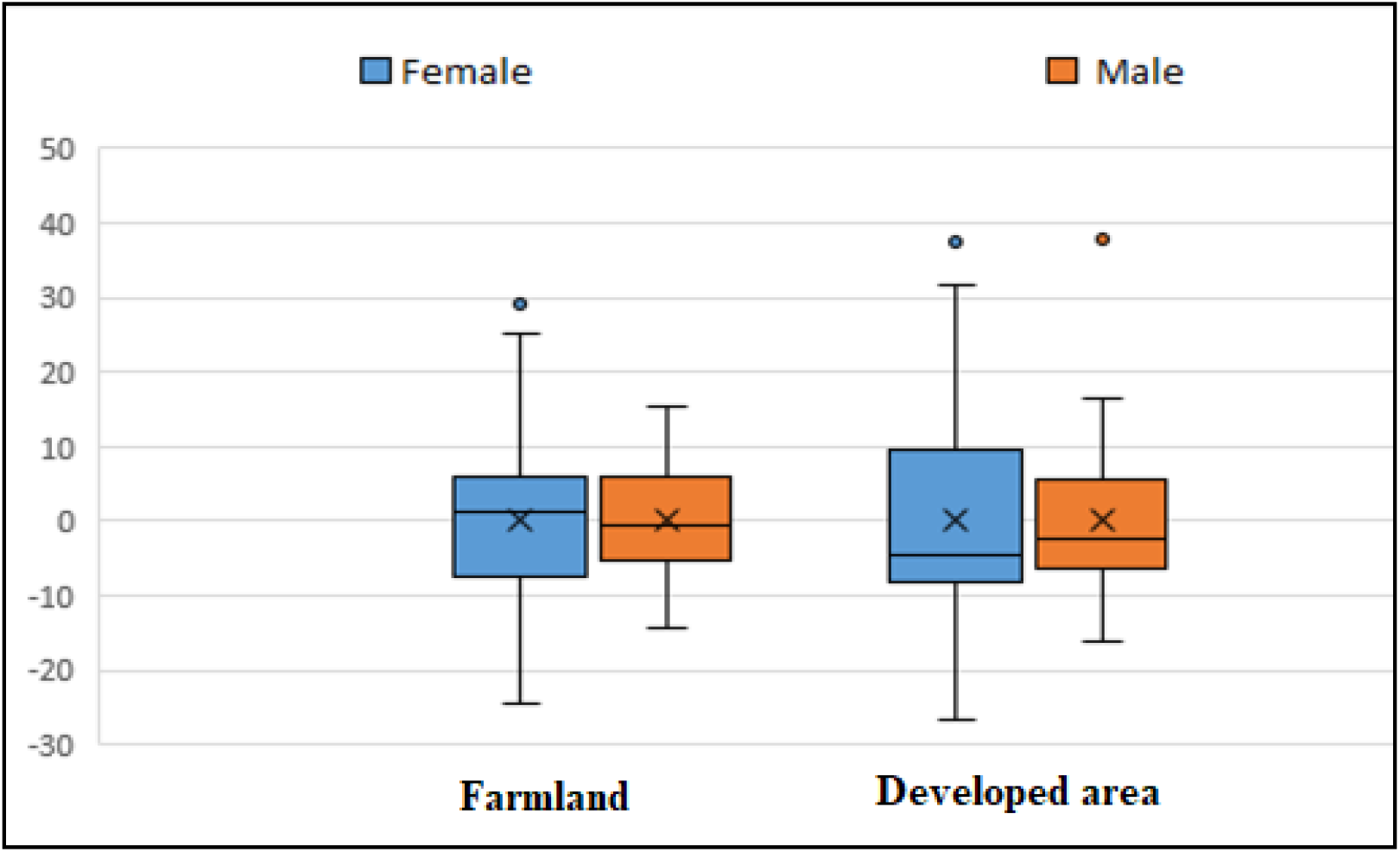
Body condition of the African common toad from the farmland and developed area

### Sexual Size Dimorphism

In the developed areas, females were significantly heavier than males (mean weights: female = 92.6 g; male = 56 g; df = 46, p < 0.00001). However, males had significantly longer (p ≤ 0.005) radio-ulna (RuL), distal forelimb (DFL), tibio-fibula (TbFL) and hind foot (FT) than females. There was no significant difference (t-test, alpha = 0.05, df = 46, p ≥ 0.345) observed in the distal hind limb (DHL), femur length (FL), humerus length (HuL), head width (HW), mouth width (MW) and snout vent length (SVL) (Table 3). Of the 21 males and 23 females were sampled at the University Farm, females showed significant higher weight, HW, and HuL (t-test, alpha = 0.05, df = 44, p ≤ 0.016). Also, females had longer FL, MW, DFL, HL, FT and SVL than males, but males had longer RuL. The differences were however not statistically significant (t-test, alpha = 0.05, df = 44, p ≥ 0.09) (Table 4).

**Table 3:**
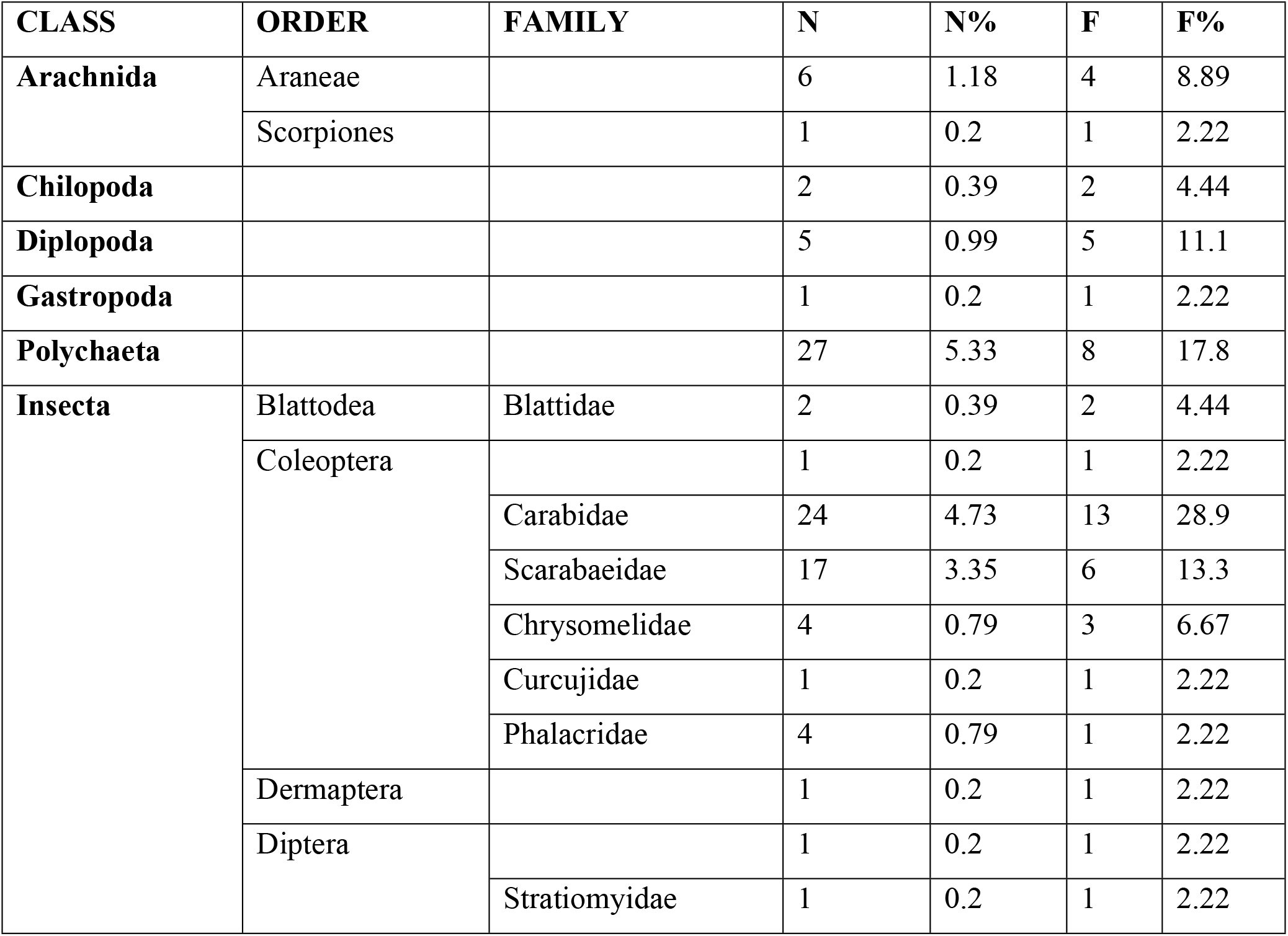
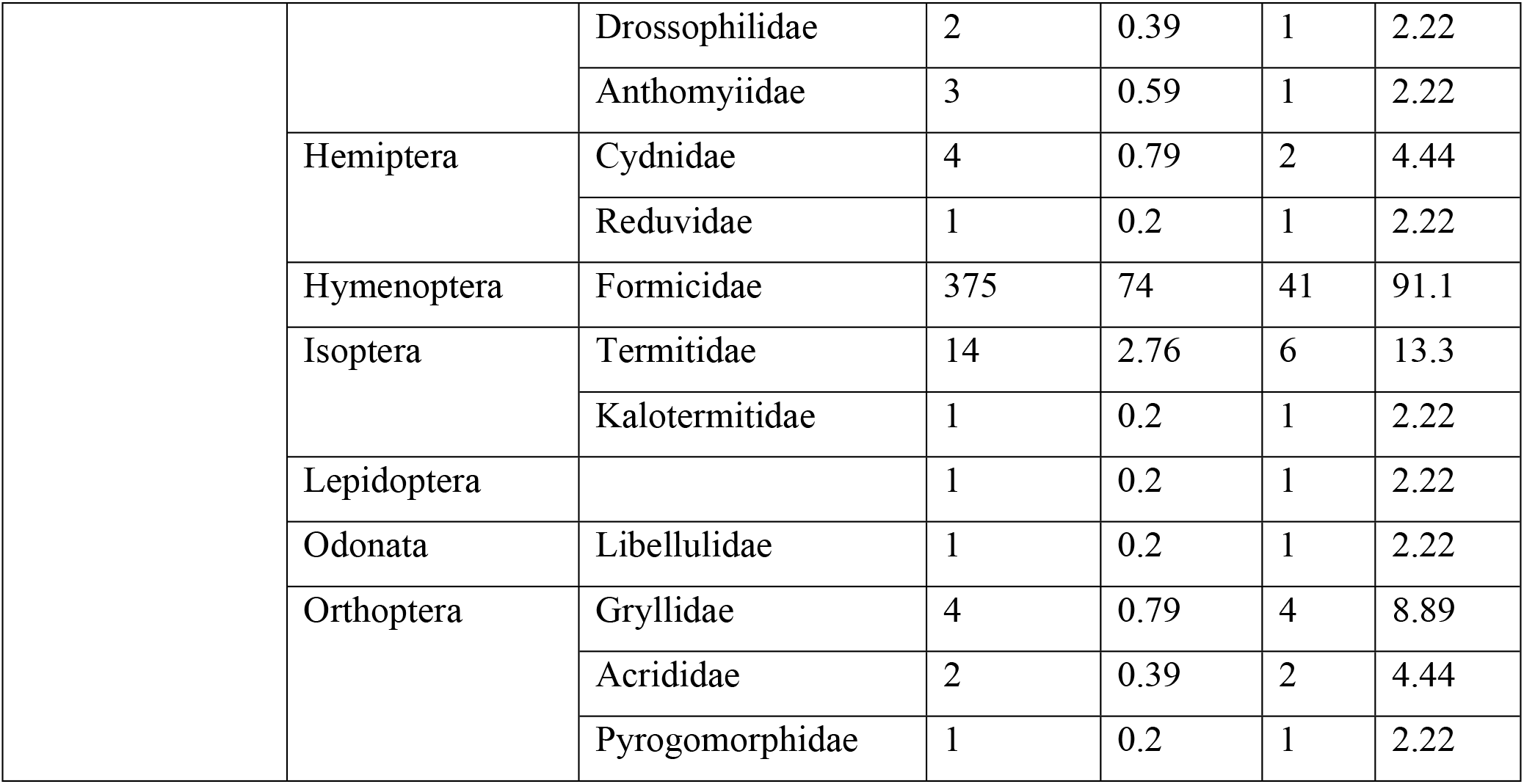
Taxonomic composition of prey items (N = 428) found in stomachs of *Amietophrynus regularis* from developed areas. (**N =** Number of items, **N% =** Numerical percentage, **F** = frequency of occurrence and **F%** = percentage frequency of occurrence).

**Table 4:** Sexual size dimorphism of African common toads from the University Farm, Legon

| Parameter | Sex | N | Minimum | Maximum | Mean | Std. Error | P-value |
| --- | --- | --- | --- | --- | --- | --- | --- |
| Distal fore limb | Male | 21 | 2 | 4 | 2.784 | 0.091775 | 0.1746 |
|  | Female | 23 | 2.2 | 4 | 2.865517 | 0.084846 |  |
| Radio-ulna | Male | 21 | 2.1 | 3.5 | 2.7 | 0.073711 | 0.9056 |
|  | Female | 23 | 2.5 | 3.8 | 2.986207 | 0.049061 |  |
| Head height | male | 21 | 2 | 2.2 | 2.157143 | 0.029738 | 0.6468 |
|  | Female | 23 | 2 | 2.5 | 2.192308 | 0.052455 |  |
| Head length | Male | 21 | 2.6 | 3.9 | 3.071429 | 0.161414 | 0.987 |
|  | Female | 23 | 2.8 | 3.3 | 3.069231 | 0.04721 |  |
| Head width | Male | 21 | 4.9 | 5.7 | 5.285714 | 0.129887 | 0.4504 |
|  | Female | 23 | 5 | 6.4 | 5.669231 | 0.110003 |  |
| Hind feet | Male | 21 | 4.1 | 4.6 | 4.285714 | 0.067006 | 0.7967 |
|  | Female | 23 | 3.7 | 4.9 | 4.323077 | 0.097503 |  |
| Weight | Male | 21 | 49 | 97 | 66.14286 | 2.656784 | 0.0161 |
|  | Female | 23 | 51 | 107 | 76.73913 | 3.225527 |  |
| Snout-vent length | Male | 21 | 10 | 13.5 | 11.2381 | 0.183935 | 0.8954 |
|  | Female | 23 | 10.3 | 20.8 | 12.06957 | 0.424796 |  |
| Distal hind limb | Male | 21 | 4.5 | 6.5 | 5.4333 | 0.08765 | 0.0431 |
|  | Female | 23 | 5.1 | 6.5 | 5.687 | 0.08419 |  |
| Femur | Male | 21 | 3 | 4.5 | 3.4143 | 0.07877 | 0.9912 |
|  | Female | 23 | 2.2 | 4.1 | 3.413 | 0.0791 |  |
| Tibio-fibula | Male | 21 | 3.1 | 5 | 3.9952 | 0.10612 | 0.5376 |
|  | Female | 23 | 3 | 5.1 | 3.9043 | 0.10069 |  |
| **Mouth width** | Male | 21 | 4 | 6 | 5.1208 | 0.09168 | 0.1806 |
|  | Female | 23 | 4 | 6 | 5.2308 | 0.1021 |  |
| **Humerus** | Male | 21 | 2.1 | 6 | 3.41 | 0.22114 | 0.1191 |
|  | Female | 23 | 2.5 | 5.8 | 3.5469 | 0.18338 |  |

**Table 5:**
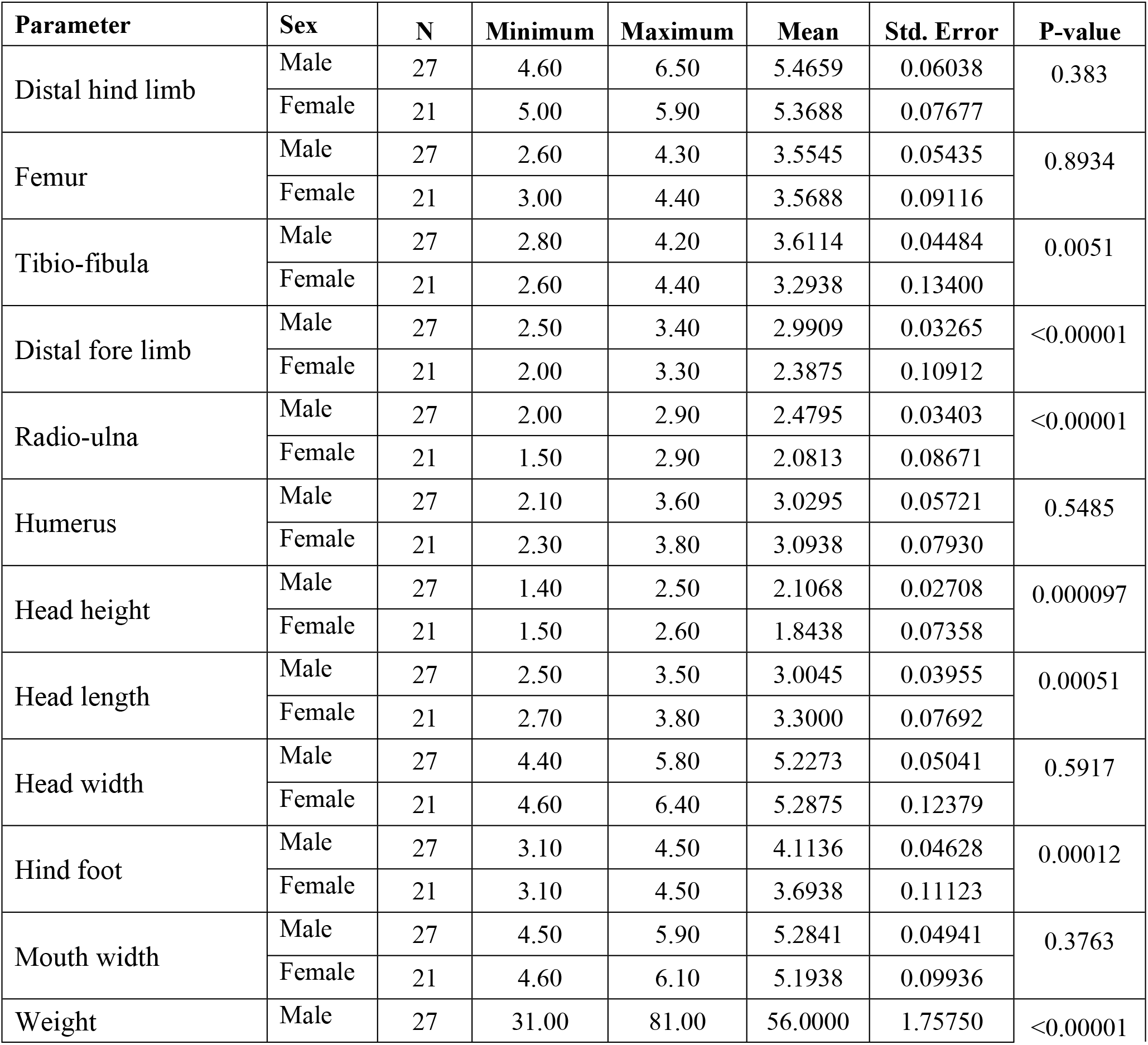
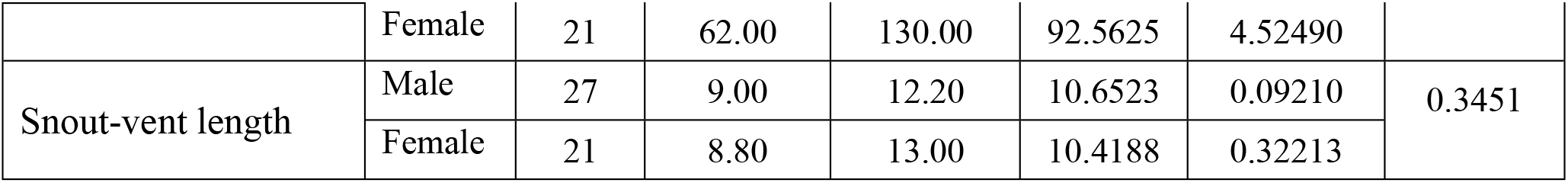
Sexual size dimorphism of toads from the Developed Area

## Discussion

### Diet Composition

Our results suggest that the common African toad has a wide taxonomic range of diet composition, and that *A. regularis* feeds predominantly on arthropods. This corroborates the results of other studies on amphibian diet, such as *Rana limnocharis* (Hirai and Matsui, 2001), *Bufo murinus* (Pamintuan and Starr, 2016), *Hyla japonica* (Park et al., 2018), *Hyla versicolor* (Mahan and Johnson, 2007), *Schoutedenella xenodactyloides* (Blackburn and Moreau, 2006), and *Scinax squalirostris* (Kittel and Sole, 2015). A high level of diet overlap (75%) observed between males and females, suggested the absence of sex-biased diet niche partitioning between the sexes. The high degree of overlap in diet composition between males and females may be due to the use of the same microhabitat for foraging and intersexual competition for food resources (Crnobrnja-Isailović *et al*., 2012).

However, females consumed a greater diversity of prey items, which could be due to their higher energy requirements, particularly during the breeding season (Quiroga *et al*., 2015), when their higher metabolic rates support reproductive functions (Balint *et al*., 2008). Also, fecundity and egg size in amphibians depend on the nutritional quality of food (Van Ngo and Ngo, 2014). Females meet high nutritional quality demands by consuming a higher diversity of prey items. Also, the diet of individual toads from the Farmland and Developed Area remained largely constant (Sorenson’s index = 83.9%; Bray-Curtis index = 91.6%), indicating the similarity in distribution of prey items between the two sites.

This was expected, given the proximity of the two sites and the fact that urbanization and agricultural practices have similar effects on invertebrate diversity and species composition (Jones and Leather, 2012). Indeed, a recent study of the ground dwelling insects in the study area showed similar species composition for the farmland and developed area, Hymenoptera (Formicidae) and Coleoptera (Carabidae) being the most abundant species (Asante, 2020). Our findings support the results obtained for *Pelophylax ridibundus* (Balint *et al*., 2008) and common toads *(Bufo bufo)* (Isailovic *et al*., 2012).

Prey items from the taxa Formicidae, Stratiomyidaea and Polychaeta were most consumed, indicating that these prey categories were the most abundant and accessible. Usually, it is the readily available and most abundant and accessible food sources that are used (Moser *et al*., 2017). The occurrence of Formicidae in the diet of most toads supports the findings of other studies on anurans (Moser *et al*., 2017; Oropeza-Sanchez *et al*., 2018). This may be due to the social behaviour and high environmental availability of these prey items (Moser *et al*., 2017). Additionally, such prey items may be relatively easy to capture and process, making their consumption energetically rewarding (Van Ngo and Ngo, 2014). The occurrence of high frequencies of Carabidae (Coleoptera) corroborates the findings for *Pelophylax ridibundus* (Balint *et al*., 2008). Common African toads are ‘sit-and-wait’ predators whose diets reflect the availability and accessibility of prey items. Their broad dietary niche therefore suggested that they are opportunistic generalist feeders.

The SVL of the toads correlated positively with prey diversity and abundance and this is consistent with the findings of Almeida *et al.* (2019). Larger individual toads have greater demand for energy uptake and so consume a higher variety and number of prey items. They also have larger stomach capacity that can accommodate large volumes of prey items compared to smaller individuals (Le *et al*., 2020). Additionally, prey consumption is influenced by activity patterns during foraging. Larger individuals are more active and tend to encounter more diverse prey items and consequently to consume more prey categories (Mageski *et al*., 2018).

### Body Condition

Body condition is a measure of energetic status, physiological state and individual quality or fitness (Cox and Calbeek, 2015; Warner *et al*., 2016). Generally, individuals in better body condition are predicted to have greater energetic status and better chance of survival compared to individuals with poorer body conditions (Labocha *et al*., 2014). In this study, body condition of individual toads was generally poor at both sites and for both males and females. This may be due to the fact that individuals experienced similar levels of food availability, accessibility, quantity and quality, and threats such as predation, competition, and parasite loads (Jarvis, 2015; Falk *et al*., 2017). Access to high quality food influences body condition and fitness, and thus plays a significant role in the success of animals (Scholz e*t al.,* 2020). The poor body condition of toads from the two study sites, Farmland and Developed Area partly suggested that the food sources are not of adequate quality. This phenomenon may be attributed to the impact of land-use/land-cover change. Even though females consumed a wider variety of prey items, their body condition was not significantly better than males. This could be due to the fact that body reserves for females might have been depleted during the breeding season (Bancila *et al*., 2010).

### Sexual Dimorphism

Our results revealed significant sexual size dimorphism in African common toads. Mating system, parental investment, diet, habitat selection, reproductive success and population density are factors that may contribute to morphological differences between male and female toads (Mori *et al*., 2017). Sexual size dimorphism may be due to different selective pressures experienced by different sexes (Lowe and Hero, 2012). For both sites, females were larger in size (SVL and weight) than males, which is consistent with the general trend in anurans and other amphibians (Lee, 2001; Monnet and Cherry, 2002; Di Cerbo and Biancardi, 2012; Lowe and Hero, 2012; Labus *et al*., 2013; Seglie *et al*., 2013; Quiroga *et al*., 2015; Mori *et al*., 2017). The disparity in weight between males and females may be due to fecundity selection (Altunisik, 2017). According to the fecundity selection hypothesis, increased fecundity of larger females is associated with increased clutch size, and consequently high reproductive success (Labus *et al*., 2013). Also, with females carrying males during amplexus, it is energetically prudent for females to be larger than males. Small size comes with agility, which makes it easier for males to reach females (Mori *et al*., 2017). It has also been reported that male amphibians usually do not live long enough to attain large sizes (Shine, 1979).

Males from the Developed Area had longer radio-ulna, distal forelimb, tibio-fibula and feet than females. The difference in tibio-fibula length is contrary to results obtained by Quiroga *et al.* (2015) on *Odontophrynus cf. barrioi*. The increase in length of such body morphologies for males may be in response to requirements for amplexus (Magalhaes *et al*., 2016). During the process of amplexus, males tend to cling onto females very tightly. This requires more muscle strength to avoid displacement by competing males. Therefore, a significant variation may be positively correlated to reproductive success in males. However, the length of humerus was similar to studies carried out by Quiroga *et al*., (2012) using *Rhinella arenarum.* According to Quiroga *et al*., (2012), this difference helps females to support their own weight and that of males on their back during amplexus. Individuals from developed areas exhibited greater levels of sexual dimorphism than those from farmland. This may have possibly been influenced by different selective regimes associated with different environmental conditions, such as intraspecific competition and predation intensity acting separately on both males and females.

### Ecological, Evolutionary and Conservation Implications

This study is probably the first to simultaneously assess the diet composition, body condition and sexual size dimorphism of *Amietophrynus regularis,* and provides vital information on the food niche and effects of human modified landscape on the body condition and sexual size dimorphism of this species. Our findings have important ecological, evolutionary and conservation implications. Individual variation in resource use have consequences for intra- and inter-specific interactions and population dynamics. Dietary patterns of individuals and populations is an essential component in trophic interactions and food web structure and provide insight into selective pressures on prey. Access to high quality food plays a significant role in the success of animals, as it influences their body condition and fitness (Scholz *et al*., 2020). Limited food availability can reduce reproductive success and survival rate, ultimately altering population densities, transient dynamics and persistence (Bolnick *et al*., 2011). The utilization of a wide range of food resources in common African toads can be advantageous in dynamic habitats with constantly changing food availability. Our results on body condition and sexual size dimorphism also provide information on how phenotypic plasticity is used by toads as an adaptation measure on an ecological time scale. In all, our data is important for understanding the feeding behaviour, population dynamics and adaptive potential of toads in changing environments. This is crucial for formulating effective conservation strategy in human-modified landscapes.

